# Early cardiac inflammation as a driver of murine model of Arrhythmogenic Cardiomyopathy

**DOI:** 10.1101/2020.06.24.169664

**Authors:** K.E. Ng, P.J. Delaney, D. Thenet, S. Murtough, C.M. Webb, E. Tsisanova, S.L.M Walker, J.D. Westaby, D.J Pennington, R. Pink, D.P. Kelsell, A. Tinker

## Abstract

The study of a desmoglein 2 murine model of arrhythmogenic cardiomyopathy revealed cardiac inflammation as a key early event leading to fibrosis. Arrhythmogenic Cardiomyopathy (AC) is an inherited heart muscle disorder leading to ventricular arrhythmias and heart failure due to abnormalities in the cardiac desmosome. We examined how loss of desmoglein 2 (*Dsg2*) in the young murine heart leads to development of AC. Cardiomyocyte apoptosis was an early cellular phenotype and RNA-Seq analysis revealed early activation of inflammatory-associated pathways in *Dsg2* null (*Dsg2*^−/−^) hearts at postnatal day 14 (Two weeks) that were absent in the fibrotic heart of adult mice (Ten weeks). This included upregulation of iRhom2/ADAM17 and its associated pro-inflammatory cytokines and receptors such as TNFα, IL6R and IL-6. Furthermore, genes linked to specific macrophage populations were upregulated. This suggests cardiomyocyte stress triggers an early immune response to clear apoptotic cells allowing tissue remodelling later on in the fibrotic heart. Our analysis at different disease stages implicate inflammation related to loss of desmoglein 2 as a major mechanism for disease progression.

## Introduction

Arrhythmogenic cardiomyopathy (AC), also referred to as arrhythmogenic right ventricular cardiomyopathy (ARVC), is an inherited heart muscle disease characterised by ventricular arrhythmias notably ventricular tachycardia and fibrillation and later in the disease process heart failure (*1, 2*). It is often a cause of cardiac arrest in young athletes. The condition appears to have acute arrhythmogenic phases before a decline in ventricular function that occurs later in the disease. Pathologically it is characterised by cardiac chamber dilatation and fibrofatty replacement of myocytes. Historically, it was thought to be a predominantly a right ventricular disease but biventricular and left ventricular patterns are now widely recognised (*1, 2*). The disease is hereditary in origin occurring in about 1 in 5,000 births and is generally autosomal dominant though recessive forms are recognised.

Both may present with prominent extracardiac features involving the skin (Palmoplantar keratoderma) and/or woolly hair. The genetic basis for the disease was first revealed by the study of autosomal recessive forms of the cardiocutaneous form of AC and demonstrated loss of function mutations in desmoplakin (Carvajal syndrome) and plakoglobin (Naxos disease) (*3, 4*). Both of these genes encode proteins that are part of the desmosome. This prompted studies in autosomal dominant AC including the examination of other desmosomal genes and revealed mutations in other components of the cardiac desmosome including desmoglein (DSG2), desmocollin2 (DSC2) and plakophilin2 (PKP2) (*5–7*). Desmosomes tether cardiomyocytes together by linking cells at the intercalated disc with the intermediate filaments, specifically desmin, in the cytoskeleton (*8*). Together with gap junctions and adherens junctions in the area composita of the intercalated disc; they provide mechanical and electrical connection between cells (*9*).

AC can be a largely asymptomatic dormant condition until the initial cardiac episode occurs in later life. These disease triggers underlying AC are still largely unknown though exercise and/or infection are proposed external factors with desmosomal dysregulation and cell adhesion as likely consequences. Early observations from myocarditis patient biopsies previously suggested the involvement of a systemic viral trigger however; no research has been able to confirm this as the cause (*10–13*). Interestingly, subsequent studies described the presence of inflammatory infiltrates in post mortem biopsies from patients with AC (*13*). Interestingly, clinical reports of myocarditis in recent paediatric cases highlight the connection between inflammation and AC (*14*). The examination of the early cardiac development in murine models of AC may provide further insight to the mechanisms that link the loss of desmoglein 2 and the inflammatory response.

There are a number of murine models for AC, including those involving genetic manipulation of *Dsg2* (*15–20*), that have been developed and studied. Global genetic deletion of *Dsg2* is embryonically lethal whilst heterozygotes with haplo-insufficiency appear normal (*17*). In contrast, transgenic mice overexpressing a missense *dsg2* mutation or a knock-in mouse lacking a domain of *Dsg2* develop cardiac dysfunction similar to that seen in AC (*15, 18*). AC can be characterised by acute episodes of worsening pathology punctuating a more stable disease course (*1*). In this study, we examine mice with the cardiac specific deletion of *Dsg2* (*19*). We describe experimental evidence and identify an early inflammatory response to cardiomyocyte apoptosis that leads to cardiac fibrosis over time.

## Results

Mice harbouring the tm1c allele for the *Dsg2* gene (Supporting Figure S1A) were crossed with the alpha MHC Cre mice (*21*) and then bred to a second generation to give the study cohort: Cre^+^, *Dsg2*^flx\flx^ (*Dsg2*^−/−^) or *Dsg2*^flx\flx^ (*Dsg2*^+/+^) mice. The results from morphometric analysis showed no differences between the adult *Dsg2*^+/+^ and *Dsg2*^−/−^ groups (Supporting Figure S1B). We also generated a small cohort of heterozygous mice (*Dsg2*^+/−^) to examine cardiac function and the effect of gene dosage in AC disease development (*20*). Most studies to date have characterised this murine model therefore, we initially focused on recapitulating this information. DAB staining, RNA-Seq and qPCR confirmed substantial decreases in transcripts translated into protein for *Dsg2* (Supporting Figure 1C). The short axis view of *Dsg2*^+/+^ and *Dsg2*^−/−^ hearts shows two morphologically different hearts at ten weeks of age. At post-mortem, the majority of *Dsg2*^−/−^ hearts showed chamber dilatation in the left ventricle and sometimes right ventricle. Some views from *Dsg2*^−/−^ hearts were also obscured due to the likely proximity of the global or segmental fibrous plaques (white arrows) which correspond to areas of fibrosis and calcinosis Cardiac assessment of left ventricular (LV) function in the *Dsg2*^−/−^ heart confirmed the LV chamber was severely dilated with little contractility. Most cardiac *Dsg2*^−/−^ mice are dead within six months, if allowed to age normally (Supporting Figure 1D).

Whilst end stage of the disease has been thoroughly explored in human AC patients and more recently with knock out murine AC models, the link between desmosome dysfunction e.g. due to loss of desmoglein 2 and the inflammatory response has not been examined in detail in these transgenic mice. We hypothesized the immune response and stress pathways in the weeks following birth may play a larger role in the immediate stages of the disease than initially reported (*19*). We studied the identity of this inflammatory infiltrate at two weeks of age and how inflammation may be associated with the progressive changes that lead to cardiac dysfunction in the adult mouse (Ten weeks).

### Adult *Dsg2*^−/−^ hearts exhibit fibrosis whereas adult *Dsg2*^+/+^ and Dsg2^+/−^ hearts appear morphologically similar

We first examined this murine model of AC by directly comparing three groups: *Dsg2* WT (*Dsg2*^+/+^), heterozygous (*Dsg2*^+/−^) and knockout (*Dsg2*^−/−^) mice at ten weeks (Figure 1). Histological analysis with i) Haematoxylin and Eosin and ii) Trichrome stain revealed prominent structural differences between *Dsg2*^+/+^ and *Dsg2*^−/−^ hearts (Figure 1A). The changes in the *Dsg2*^−/−^ heart were pronounced at adulthood with evidence of myocyte death and interstitial fibrosis i), collagen deposits ii) and in some severe cases, calcification which appear as dark purple deposits within the fibrotic area. No abnormal morphological changes or signs of fibrosis were observed in the *Dsg2*^+/+^ and *Dsg2*^+/−^ hearts.

**Figure 1:**
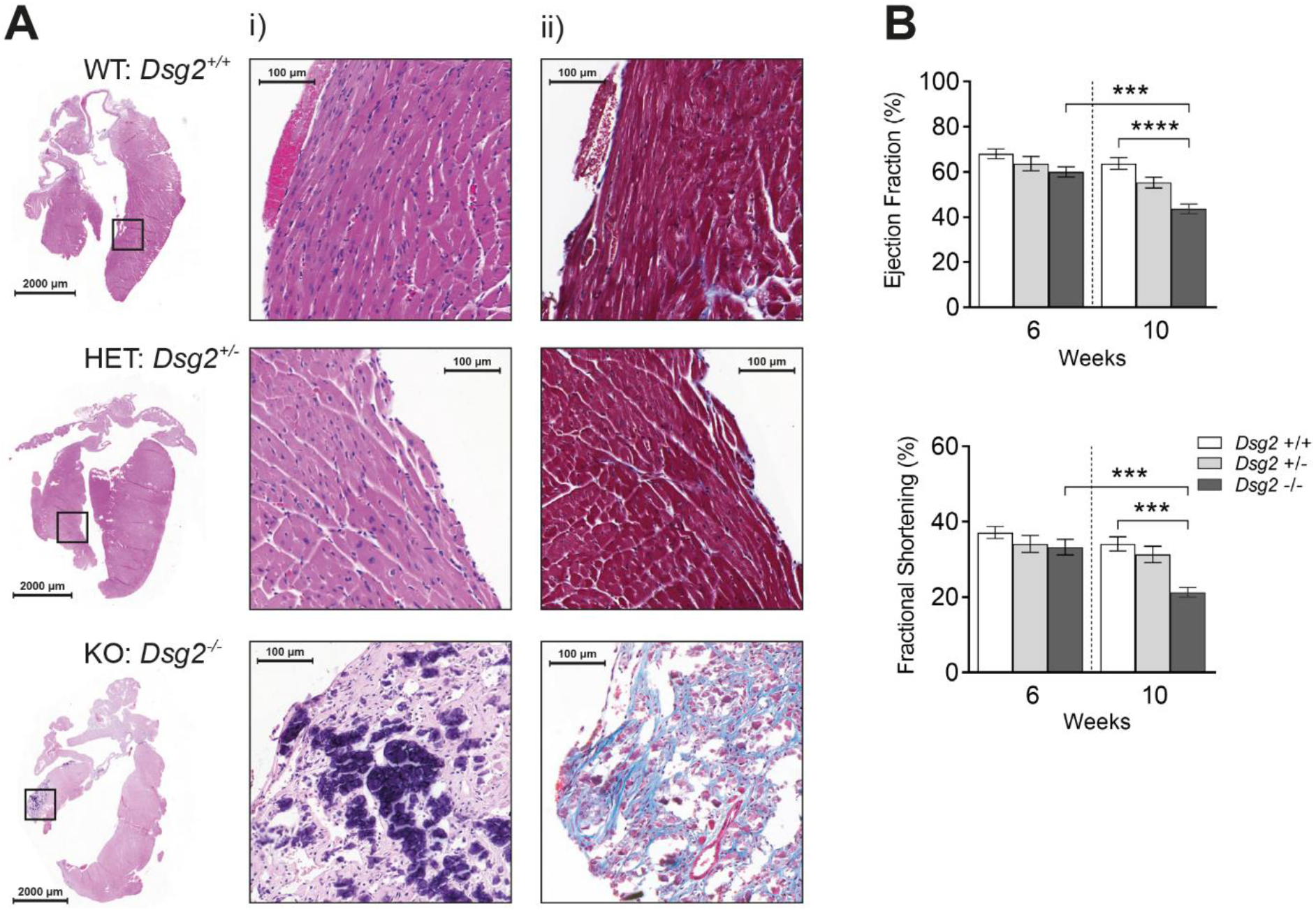
Histological and cardiac assessment of cardiac specific desmoglein 2 mouse model. The Dsg2^−/−^ adult mouse exhibits severe cardiac dysfunction due to gross morphological defects. (A) Detailed histology of whole hearts at ten weeks with zoomed in areas: i) H and E and ii) Trichrome equivalent in wild-type (WT) *Dsg2*^+/+^, heterozygous (HET) *Dsg2*^+/−^ and knockout (KO) *Dsg2*^−/−^ hearts. The *Dsg2*^+/+^ and *Dsg2*^+/−^ hearts both appear normal at ten weeks with no abnormal morphology. The extreme phenotype in the *Dsg2*^−/−^ heart exhibits changes within the myocardium with extensive calcinosis (Purple deposits in H and E and fibrosis (Blue stain for collagen in Trichrome stain). (B) Cardiac assessments of this mouse model showed there were no significant differences in cardiac function at six weeks between all groups (n= 8, all groups). However, ejection fraction (%) and fractional shortening (%) was significantly reduced between *Dsg2*^+/+^ and *Dsg2*^−/−^ hearts by ten weeks ***p < 0.001 and *** p<0.0001. There is also a significant decline in cardiac function in *Dsg2*^−/−^ heart as the mice age from six to ten weeks ***p < 0.001. All graphs represent Mean ± SEM; data was analysed by one-way ANOVA with Tukey's multiple comparisons test.

The echocardiographic assessments from the three genotypes are shown in Figure 1B. We measured cardiac function in younger juvenile mice (Six weeks) and adult mice (Ten weeks). The heterozygous group were also included in the analysis to reveal if the decline in cardiac function is related to gene dosage (n=8, all groups). At six weeks of age, there are no statistically significant differences however, the severe morphological changes in the diseased heart at ten weeks result in the substantial decline in ejection fraction (**** p<0.0001) and fractional shortening (*** p<0.001) indicating functional impairment of contractility (Figure 1B). The *Dsg2*^−/−^ mice worsen with age whereas the *Dsg2*^+/+^ mice exhibited normal cardiac function. The *Dsg2*^+/−^ mice also show a slight decrease in cardiac function.

We examined the electrophysiological properties of the heart using electrocardiogram (ECG) traces from cardiac echocardiographic assessments acquired at ten weeks. The ECG parameters of interest were compared between *Dsg2*^+/+^ and *Dsg2*^−/−^ cohorts (n=8, both groups). Heart rates were normal (Beats per minute (BPM) and RR interval), there were no differences with PR interval, P wave duration and QRS interval however there was a significant increase in QT and QTc intervals, * *p* < 0.05 in the cardiac *Dsg2*^−/−^ mice (Supporting Figure 2A). We further investigated the electrophysiological substrate in Langendorff-perfused isolated hearts using FlexMEA array as previously described (*22, 23*). S1S2 decremental protocols were performed and conduction velocity, mean increase in delay, a sensitive measure of conduction delay and effective refractory period were measured. Capture was however; inconsistent and challenging with *Dsg2*^−/−^ hearts, as the fibrotic plaques were difficult to avoid however, there were no significant changes in these parameters (Supporting Figure 2B).

### Loss of cardiac Dsg2 at two weeks present with signs of inflammation

Our studies at ten weeks show a highly fibrotic and failing heart. The question arises as to what are the initial disease processes that promote this pathological state. To address this, we characterised mice at two weeks of age (Figure 2). The majority of morphological observations from two week old *Dsg2*^−/−^ hearts showed distinctive small surface lesions. The comparison with the pathology in the ten week hearts is interesting: the plaque-like lesions were larger consistent with fibrosis (Figure 2A). Many lesions were predominately present on the right epicardial surface although a large number were also located within intraventricular septum and were not present in littermate controls (Figure 2B). Detailed histological analysis revealed the lesions contain densely cellular infiltrates. These pronounced areas of cellular infiltrates (black arrow) are present in the i) H and E sample with the white plaques surrounded by existing cardiomyocytes (blue arrow). On closer inspection, ii) Trichrome stain show the lesions present with early subtle changes in the extracellular matrix leading to interstitial fibrosis. Our findings suggest that loss of cardiac *Dsg2* at this postnatal stage leads to inflammation that precedes fibrosis in the adult *Dsg2* null heart. There were no abnormal areas or signs of inflammation in two week old *Dsg2*^+/+^ hearts.

**Figure 2:**
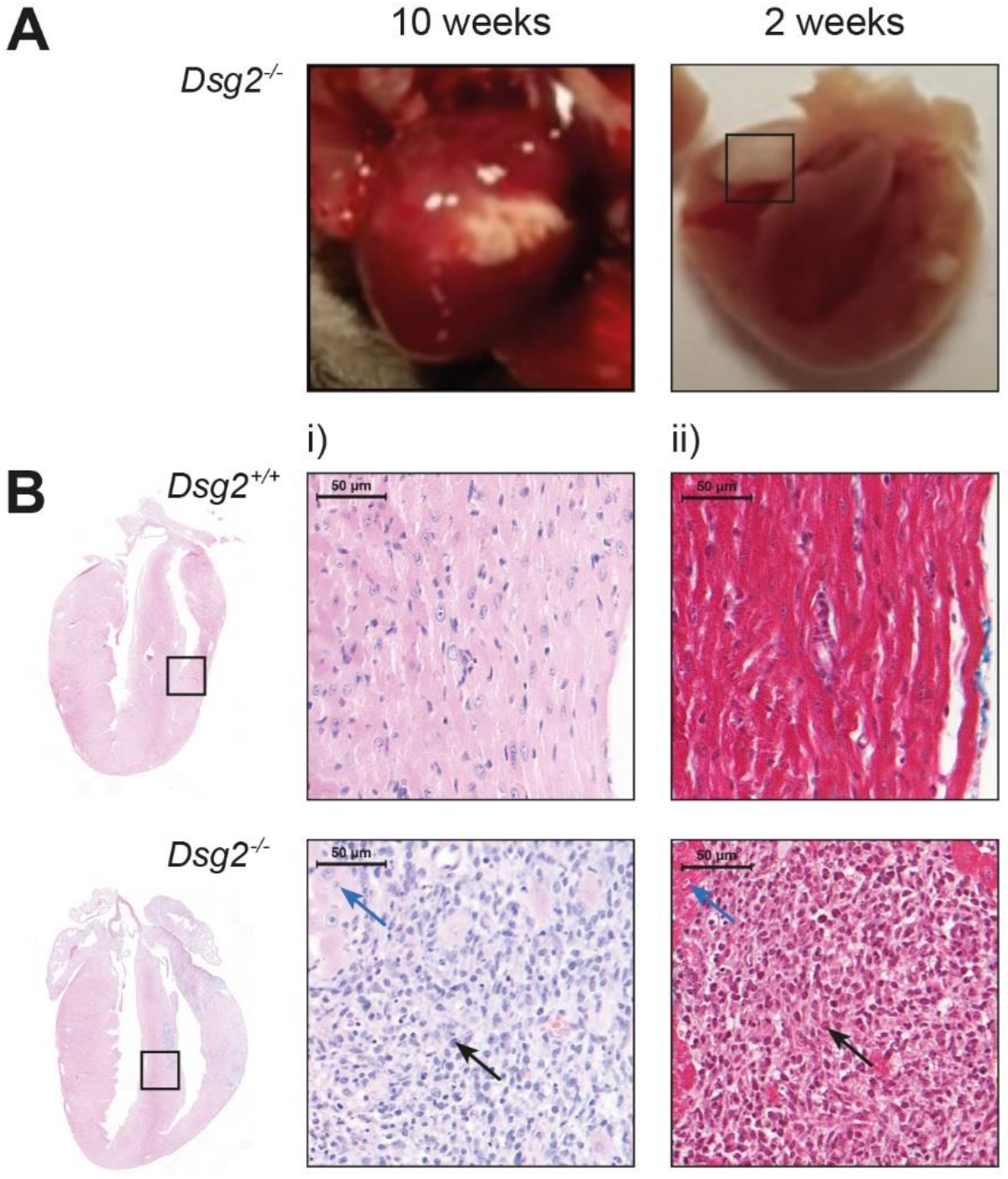
Gross morphological changes are evident in *Dsg2*^−/−^ hearts at two weeks of age. Histological analysis confirms early signs of inflammation. (A) Gross morphological comparison of the adult (ten week) and postnatal (2 week) *Dsg2*^−/−^ heart. Large dense fibrous plaques are present in the adult heart where changes in the extracellular matrix have already taken place. We also observed the presence of much smaller plaques in the postnatal heart. (B) Detailed histology of *Dsg2*^+/+^ and *Dsg2*^−/−^ hearts at two weeks: i) H and E and ii) Trichrome stain. The myocardium in *Dsg2*^+/+^ heart is normal with healthy myocytes. However, the H and E analysis for the postnatal *Dsg2*^−/−^ heart presents extreme phenotype and inflammation with the accumulation of dense nuclei, predominantly immune cells (black arrow) and some normal cardiomyocytes (blue arrow). The *Dsg2*^−/−^ hearts do not stain positive for collagen (ii) but there are clear changes within these plaque like areas within the myocardium at this early stage of development.

### Comparison of cardiac gene expression at two and ten weeks

We next performed and examined bulk RNA-Seq data from mouse heart tissue from two and ten week cohorts to identify potential pathways that may be involved in the early pathogenesis of this disease. There are a number of significantly differentially expressed genes at two weeks when compared to the ten weeks data set with some overlap of common genes between the postnatal and adult period (Figure 3A). KEGG pathway analysis revealed striking differences between the two time points with quite distinct pathways that are upregulated at two weeks. These include signalling pathways linked to cell regulation and inflammatory like Janus kinases (JAKs), signal transducer and activator of transcription proteins (STATs), growth factors and pro inflammatory cytokines such as Interleukin 6 (IL-6) (Figure. 3B). In contrast, the KEGG pathway analysis at ten weeks revealed different upregulated pathways, which are linked to changes in extracellular matrix and tissue remodelling (Supporting Figure 2C). It has been well documented that fibrosis is evident and the result of desmosome dysfunction in this murine model of AC (*19, 24*). We explored if the failing hearts in the young mice start activating fibrotic pathways linked to tissue remodelling and scar formation, we measured the expression of fibrosis markers *Col1a1*, *Col3a1*, *Ctgf, Tgfβ1* and *Tgfβ2* at two and ten weeks with qRT-PCR. *Col1a1*, *Col3a1*, *Ctgf* and *Tgfβ1* markers were significantly upregulated at two weeks in *Dsg2*^−/−^ hearts, although *Tgfβ2* did not change in the early stages of the disease (Supporting Figure 2D). At ten weeks, *Cola1*, *Col3A1* and *Tgfβ1* expression all declined at ten weeks, *Ctgf* was highly expressed at both time points whilst *Tgfβ2* increased significantly in the adult *Dsg2*^−/−^ heart.

**Figure 3:**
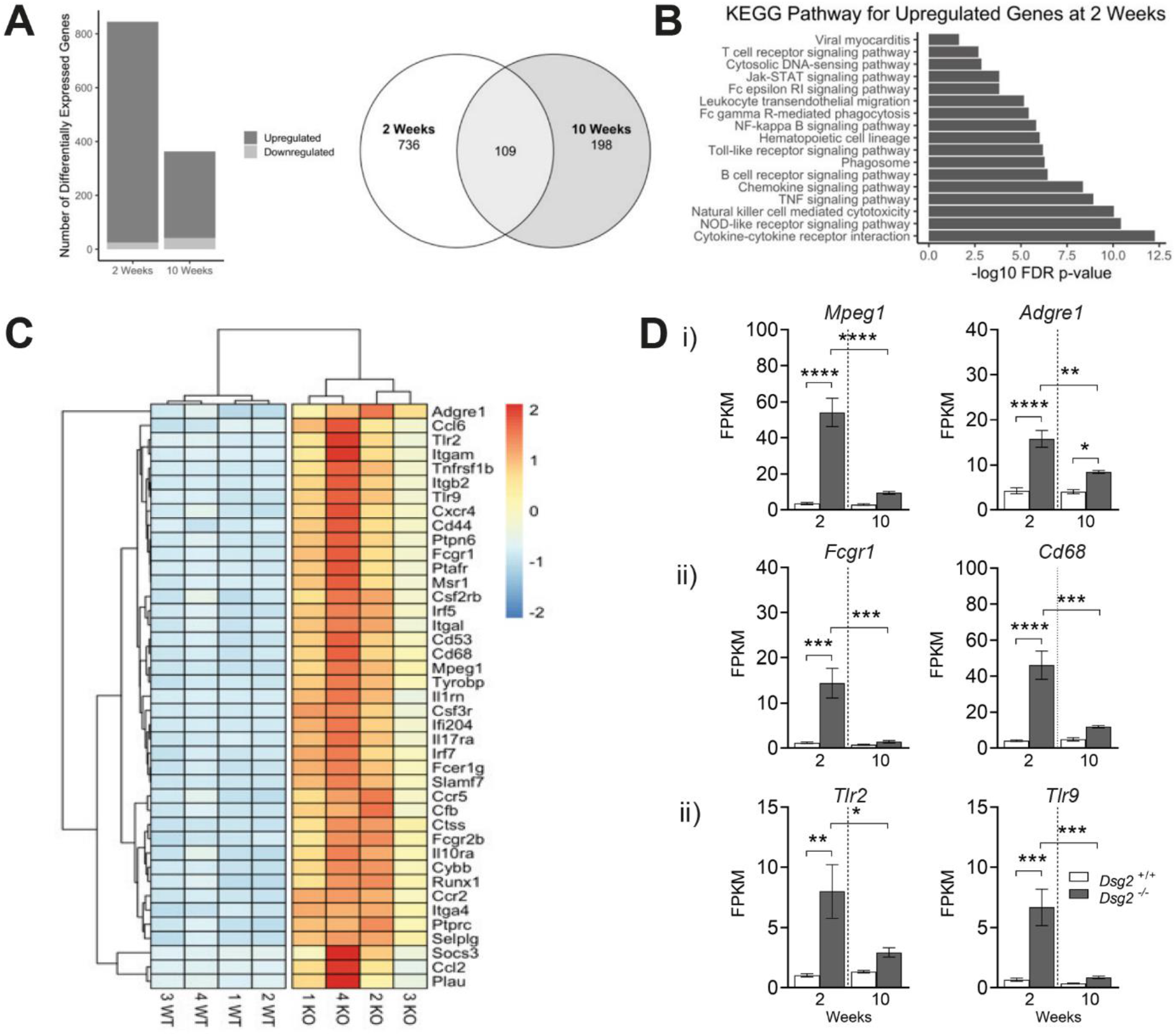
RNA-Seq reveal specific immune cell populations such as macrophages are upregulated in *Dsg2*^−/−^ hearts at two weeks of age. (A) Bar plot depicting the number of differentially expressed genes, both upregulated and downregulated, between *Dsg2*^+/+^ and *Dsg2*^−/−^ mice at two and ten weeks (*Dsg2*^+/+^ and *Dsg2*^−/−^ n= 4, for both timepoints). The Venn diagram represents the number of differentially expressed genes between *Dsg2*^+/+^ and *Dsg2*^−/−^ groups at the two timepoints and the number of genes which crossover between both groups. (B) KEGG Pathway analyses for upregulated genes at two weeks (C) Heatmap was generated from 41 differentially expressed immune-related genes. (D) Transcriptome data (FPKM = Fragments Per Kilobase of transcript per Million mapped reads) show macrophages are the main inflammatory cell population present in *Dsg2*^−/−^ hearts. Genes of interest include: (i) *Adgre1* (F480) and *Mpeg1*, both highly expressed in mature macrophages (ii) *Fcgr1* and *Cd68*, are expressed in monocytes and macrophages and (iii) *Tlr2* and *Tlr9.* These genes trigger the innate immune response. Graphs represent Mean ± SEM, where *p < 0.05, \*\**p* < 0.01, \*\*\**p* < 0.001, \*\*\*\**p* < 0.0001.

The Heatmap was generated from 41 differentially expressed immune related genes at two weeks (Figure 2C). The genes of interest include a significant increase in the inflammatory cell marker *CD45 (Ptprc*) which support the histological observations of inflammatory infiltrates at the lesion areas in the postnatal *Dsg2*^−/−^ heart. In depth analysis of the transcriptome data revealed the main population cells in the lesion may be predominately macrophages (Figure 3D). Evidence for this include i) Activated and mature macrophages express specific genes: Adhesion G protein-coupled receptor E1 (*Adgre1, or its common name F4/80)* and macrophage expressed gene 1 *Mpeg1*. Macrophages can polarise into the different subsets with distinct functional roles ii), the transcriptome data suggest the two week old *Dsg2*^−/−^ hearts include M1 classically activated macrophages (M1 expressing *Fcgr1* and *Cd68*) (*25, 26*). Cells of monocyte lineage express CD68, a protein highly expressed by monocytic phagocytes, macrophages and osteoclasts. This gene is also expressed by circulating and tissue macrophages. Toll-like receptor protein 2 (*Tlr2* or *Cd282*) and *Tlr9 (Cd289)* (iii) both recognise foreign material and can initiate the innate immune response. We report M2 alternatively activated macrophages were also present, mouse specific M2 cell surface markers such as *Fizz1*, *Ym1/2* and *Arg1* were detected at two weeks of age (*27*). Please refer to Table 1 for other macrophage genes upregulated in the two week *Dsg2*^−/−^ mouse heart. Thus, these receptors are expressed by a wide range of inflammatory cells, these include macrophages although the prolonged activation of these receptors may also be detrimental (*28*). The expression of these immune associated genes decreased by ten weeks with no differences between *Dsg2*^+/+^ and *Dsg2*^−/−^ hearts at adulthood, this was previously noted in the ten week KEGG pathway analysis (Supporting Figure 2C).

**Table 1:**
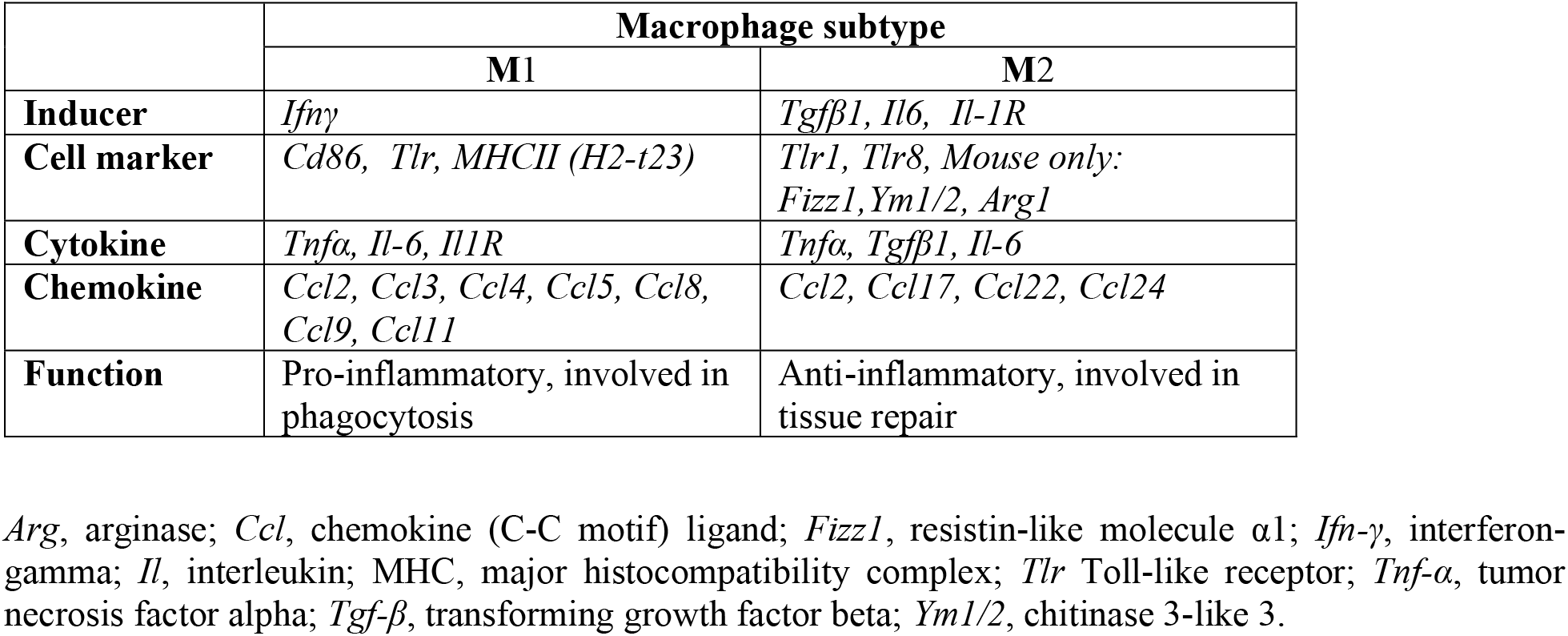
Phenotype and function of macrophage subsets identified with our transcriptome data set (Two weeks versus Ten weeks). Detailed analyses reveal specific macrophage genes were significantly upregulated in the Dsg2^−/−^ mouse heart at two weeks.

### The inflammatory infiltrate observed at two weeks consists of macrophages

The postnatal period of cardiac growth is clearly an important stage of development in murine *Dsg2*^−/−^ hearts. Histology and RNA-Seq data suggests inflammation may be a key driver in development of AC in this mouse model. We further characterised the cellular infiltrate observed in lesion areas with DAB chromogenic staining for inflammatory cells with CD45 and F4/80 for macrophages (Figure 4A). Littermate control (*Dsg2*^+/+^) hearts showed no inflammatory cells within the myocardium with the majority located within the blood vessel. However, in the myocardium of cardiac specific *Dsg2*^−/−^ mice, there was an increase in inflammatory cell marker CD45 in the lesional areas previously described in Figure 2A and B. The inflammatory cells accumulate within the 1) intraventricular septum and 2) epicardial surface (Figure 4A). The DAB staining also confirmed the presence of macrophages (F4/80) in the *Dsg2*^−/−^ heart. The remainder of CD45 + cells may also consist of monocytes, granulocytes and natural killer cells. The increase in CD45+ cells in two week old *Dsg2*^−/−^ hearts when compared to adult hearts (Ten weeks) was confirmed by i) qPCR analysis and ii) RNA-Seq (Figure 4B). The presence of macrophage recruitment is further supported by evidence that the chemokine monocyte chemotactic protein (MCP-1, gene *Ccl2*) is also highly expressed (Figure 4C). Its receptor C-C Motif Chemokine receptor 2 (*Ccr2*) binds to MCP-1 and allows activated macrophages to initiate the inflammatory response (*25*). These genes are active during the innate immune response that controls the migration and infiltration of monocytes/macrophages.

**Figure 4:**
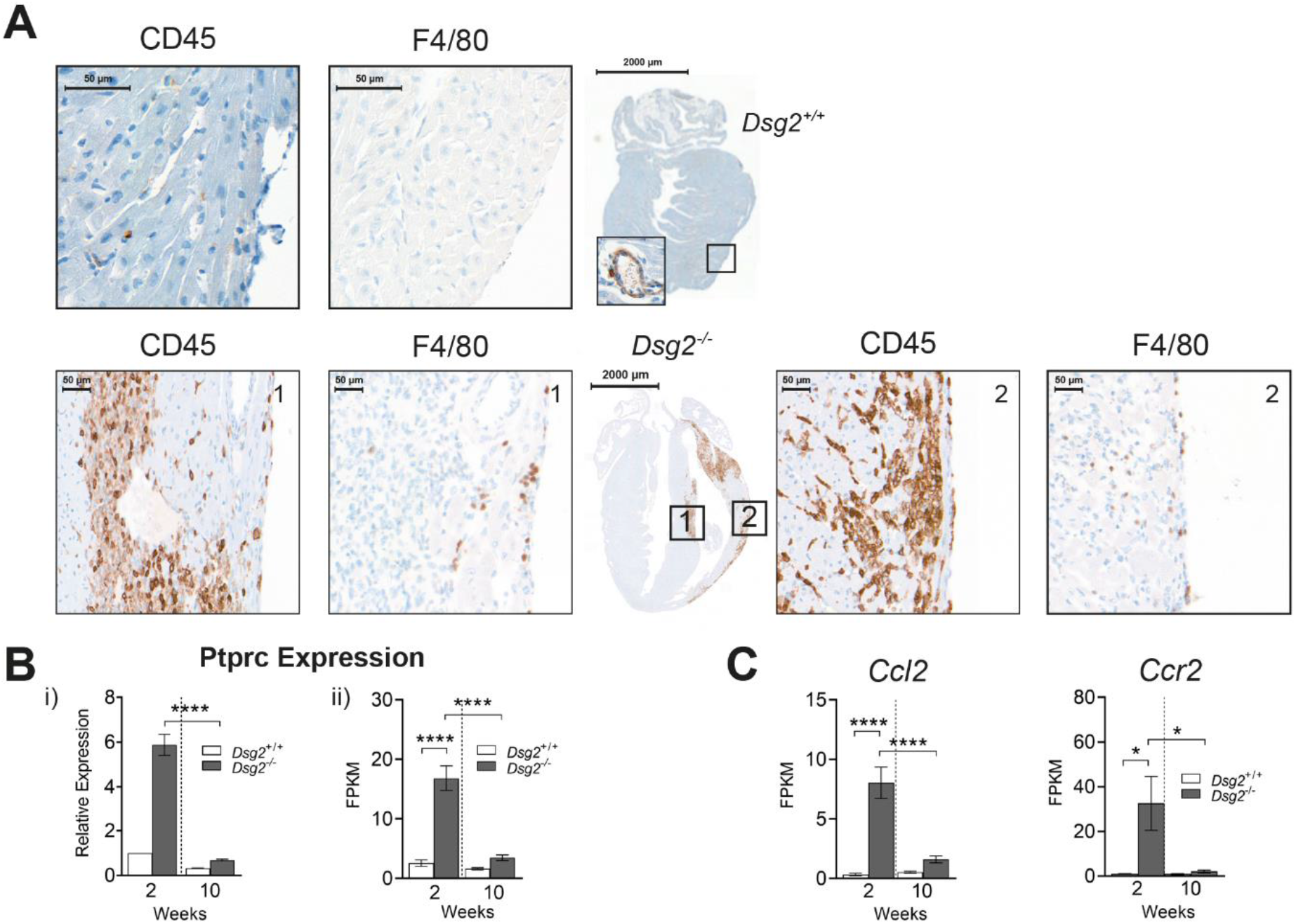
Inflammation and cells from the innate immune response play an important role in the early stages (two weeks) of desmoglein 2 murine AC disease progression. (A) DAB immunostaining analysis in two week old *Dsg2*^+/+^ and *Dsg2*^−/−^ hearts. The zoomed in areas for each heart show the same region where we have stained for CD45 and F4/80. There are no immune cells present within the myocardium of the *Dsg2*^+/+^ heart; however, a large population of CD45+ cells are present in the large vessels of the heart. The *Dsg2*^−/−^ heart exhibits increased expression of CD45+ cells within the 1) intraventricular septum region of the myocardium 2) and epicardium; these areas of inflammation include some macrophages. (B) To validate the CD45+ DAB results, we performed i) qRT-PCR (n = 5, all groups RE = Relative expression) and ii) RNA-Seq (n=4, all groups) confirming the presence of CD45+ cells (*Ptprc*) in *Dsg2*^−/−^ hearts at two weeks of age when compared to their *Dsg2*^+/+^ littermates. Data represent Mean ± SEM, \*\*\*\**p* < 0.0001. These results confirm this unique CD45+ signature is only observed in Dsg2^−/−^ postnatal hearts. (C) Transcriptome data to validate the activation and recruitment of macrophages in two week old *Dsg2*^−/−^ hearts. *Ccl2*, its receptor *Ccr2* were both highly expressed during this inflammatory stage (n= 4, for all groups). All data represent Mean ± SEM, *p<0.05 and \*\*\*\**p* < 0.0001.

To shed more insight into these results, we further examined the inflammatory infiltrate observed at two weeks with FACS analysis. Specific gating strategies were employed to isolate and identify the immune cell population from whole hearts. Cells were isolated with CD45, CD11b and F4/80 antibodies (n=7, both groups) (Figure 5A). CD45 is an inflammatory cell marker, present on all hematopoietic cells apart from erythrocytes and plasma cells. There was a significant increase in CD45+ cells in the *Dsg2*^−/−^ heart when compared to their littermate controls and specifically large CD11b positive population that include cells of the innate immune response such as neutrophils, monocytes, granulocytes and macrophages. Earlier transcriptome analysis suggested the presence of macrophage in the young diseased heart, therefore we gated with F4/80 which confirmed a high proportion of these CD45+ cells were macrophages. At this young age, it is expected that the adaptive immune response would not play a significant role in these animals, yet it was of interest to examine whether the T cell population were present in the large population of CD45+ cells we identified with i) CD3 marker and other T cell subsets such as i) T cytotoxic and T helper cells with CD4 and CD8 antibodies respectively (Figure 5B). The number of T cells isolated from these mice was relatively low in both groups suggesting that the adaptive immune response does not have a key role during the early stages of murine AC disease pathogenesis. From previous DAB, RNA-Seq and histology results and further experimental evidence from ii) FACs analysis in Figure 5B also suggests that the large population of CD45+ cells are macrophages in theses *Dsg2*^−/−^ hearts when compared to their littermate controls.

**Figure 5:**
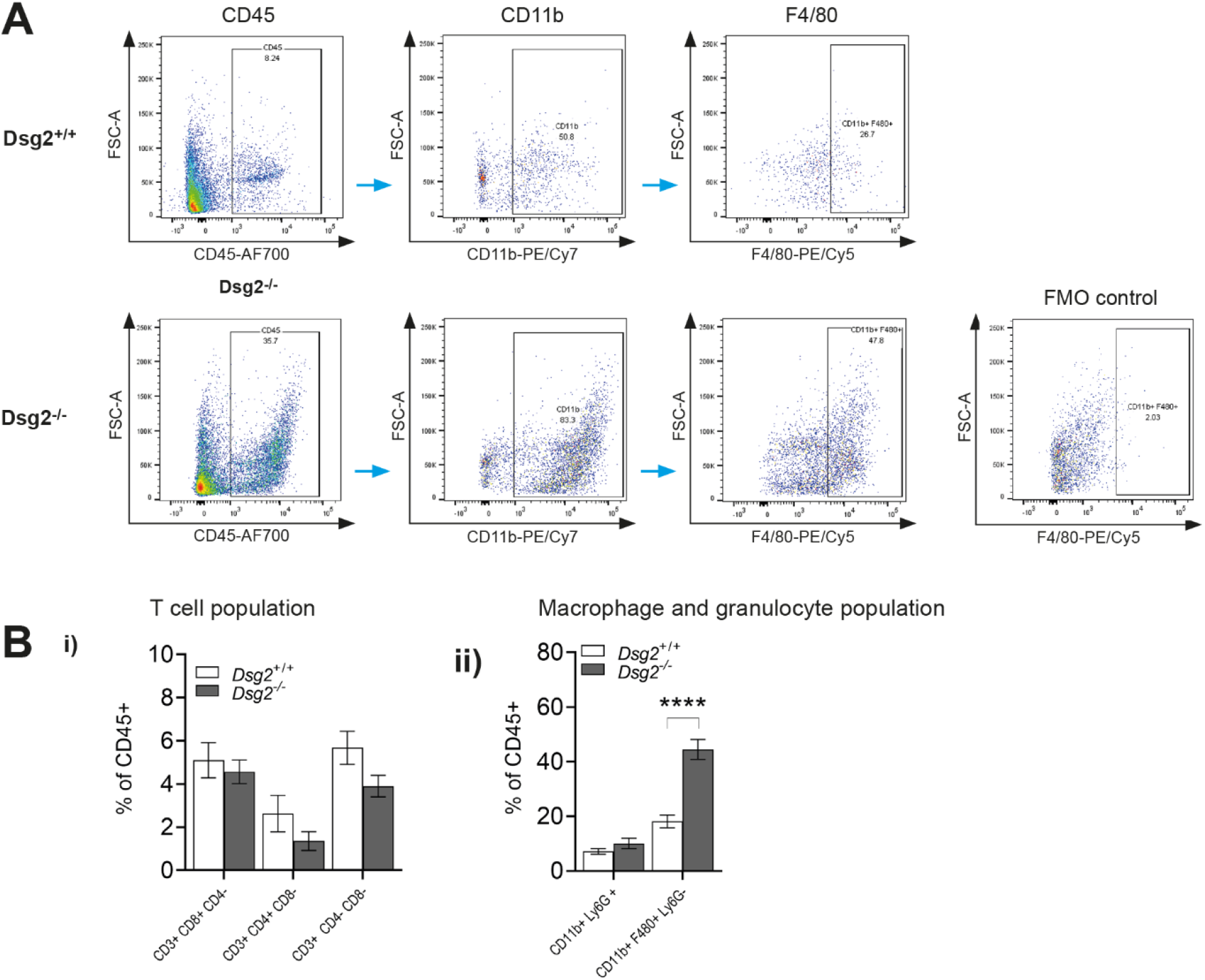
The unique inflammatory signature observed in the *Dsg2*^−/−^ postnatal heart (Two weeks) contains a large macrophage population. (A) Example plots of the gating strategies to identify macrophages with CD45, CD11b and F4/80 antibodies in two week old *Dsg2*^+/+^ and *Dsg2*^−/−^ hearts. The *Dsg2* null heart contains a large population of CD45+ cells that are predominately macrophages (B) Detailed FACs analysis of CD45+ cell population at two weeks in *Dsg2*^+/+^ and *Dsg2*^−/−^ hearts (n= 7, for both groups). Bar graphs to show the distribution of CD45+ cells, cells were either i) CD3+ T cells populations or CD3-giving ii) Macrophage or granulocyte population. There is a large macrophage population in *Dsg2*^−/−^ hearts when compared to their littermate controls. Data represent Mean ± SEM, \*\*\*\**p*< 0.0001.

### The activation of specific stress pathways occurs alongside cardiomyocyte death

In a normal situation, macrophages and cells of the innate immune response are activated and recruited to sites of inflammation when tissue injury or infection is detected by the organism (*28*). Macrophages have a wide range of functions; they are phagocytic and are recruited by specific intracellular signals to remove infected/dying cells or they may be involved in tissue repair (*26*). We performed TUNEL analysis on the two week samples and showed an increased number of myocytes were undergoing apoptosis; 8 % of cells in *Dsg2*^−/−^ heart compared to 0.47 % in the *Dsg2*^+/+^ heart (Figure 6A and 6B). The RNA-Seq data set also revealed mitochondrial apoptosis inducing factor 1 (*Aifm1*) was also upregulated at two but not ten weeks (Figure 6C). Our results suggest stressed cardiomyocytes that undergo apoptosis in these hearts may initiate specific signalling pathways that trigger the early inflammatory response observed.

**Figure 6:**
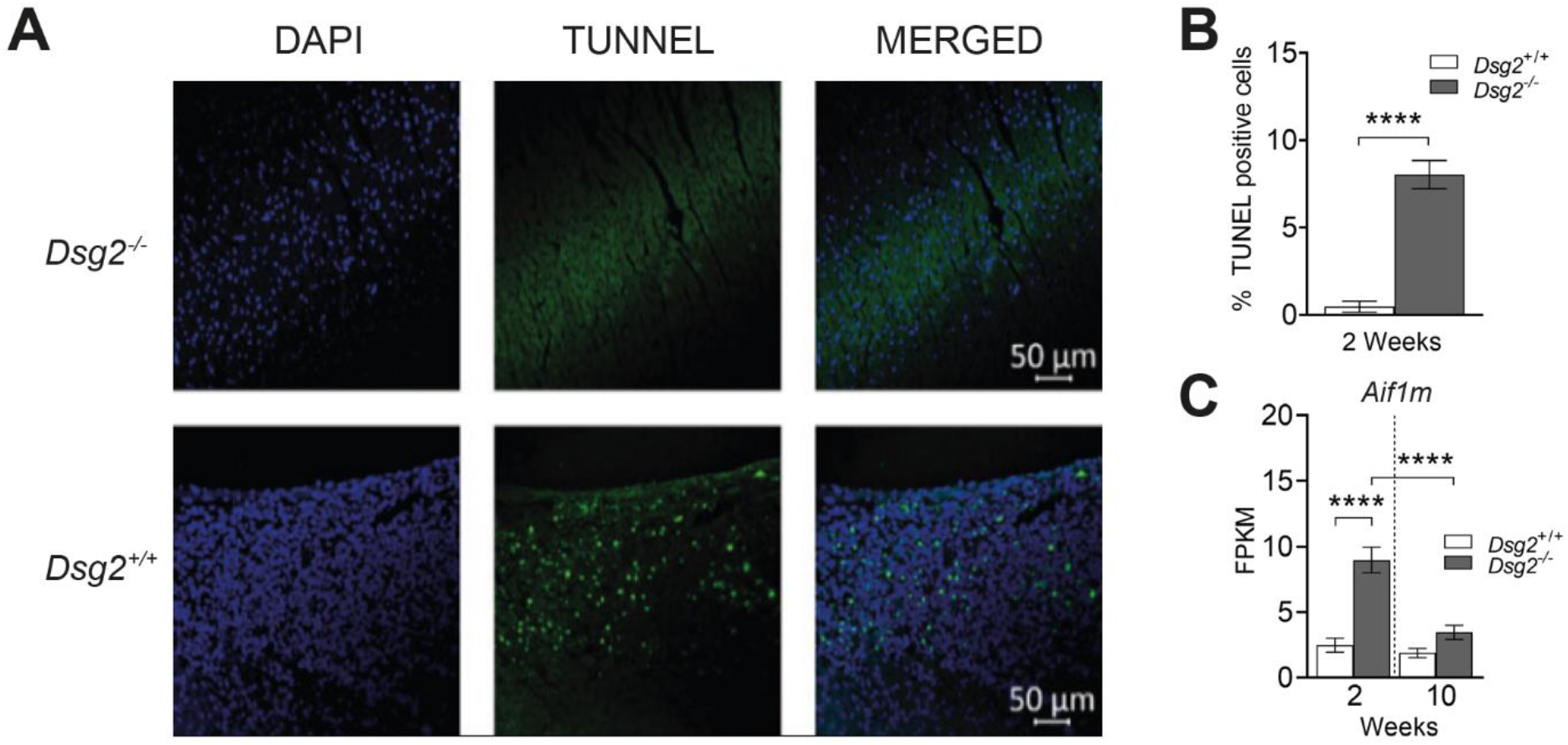
Apoptosis and inflammation occur simultaneously in the diseased postnatal heart. (A) TUNEL analysis in two week old *D*sg2^+/+^ and *Dsg2*^−/−^ hearts. From earlier histological observations, the areas where there is interstitial fibrosis, there is also apoptosis in the *Dsg2* null heart. (B) Bar graph to show there is a significant number of TUNEL positive (apoptotic) cells in the *Dsg2*^−/−^ hearts when compared to their control littermates (n= 3, for both groups). (C) RNA-Seq analysis reveals that the mitochondrial gene, apoptosis-inducing factor 1 (*Aif1m*) is also upregulated at two weeks suggesting nuclear disassembly during apoptosis (n= 4, for both groups). All data represent Mean ± SEM. \*\*\*\**p* < 0.0001.

One pathway of interest with a major inflammatory function is the iRhom2/ADAM17 stress pathway. The seven-membrane spanning protein iRhom2 (inactive Rhomboid 2 encoded by *Rhbdf2*) was found to regulate and promote maturation of ADAM17 (*29*). Mature Adam17 promotes the cleavage of a number of ligands such as Il-6r, Tnfα and *Dsg2*. Cardiac stress can activate the IL-6 family of receptors, which include profibrotic IL-11 (*30*). On closer inspection, Il-6r was highly expressed in the Dsg2^−/−^ heart at two weeks (Figure 7A). Results from i) qRT-PCR and ii) RNA-Seq confirm that iRhom2 and Adam17 were both upregulated at two but not ten weeks. In our hands, the levels of Il-6 were too low to detect with our RNA-Seq data set however, we were able to detect an increase in Il-6 from i) qRT-PCR and Il-6r ii) transcriptome data. This surge of Il-6 may be released directly from macrophages and\or stressed cardiomyocytes, facilitating changes in the myocardium from a physiological to pathological state. Thus, these results highlights how iRhom2/Adam17 stress pathway may initiate the innate immune response considering its diverse roles in growth factor signalling and regulation of protein maturation and secretion (*31*). Additional cytokines of interest include Tumor necrosis factor alpha (Tnfα) and Transforming growth factor beta 1 (Tgfβ1), which are elevated at two weeks in the *Dsg2*^−/−^ mouse heart (Figure 7B). These upregulated genes facilitate further inflammation, apoptosis and changes in extracellular matrix.

**Figure 7:**
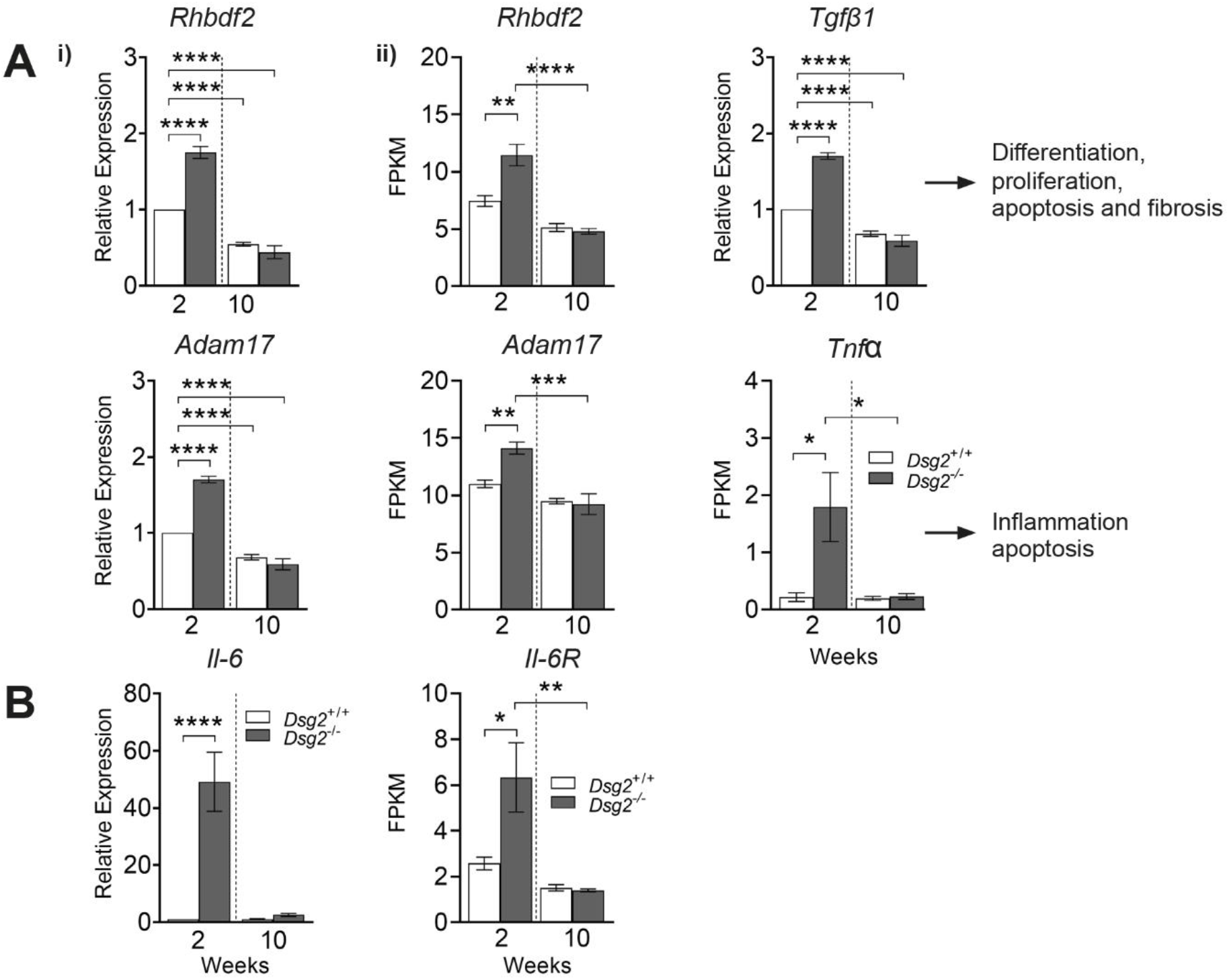
Transcriptome data reveals early activation of stress and inflammatory-associated pathways in two week old *Dsg2*^−/−^ hearts when compared to ten week old adult cohort. (A) The iRhom2/ADAM17 stress pathway is upregulated in the two week old *Dsg2*^−/−^ heart. Quantitative PCR analysis (i) and RNA-Seq (ii) confirms the activation of *Rhbdf2* and *Adam17* at this early time point (qRT-PCR n= 5, for both groups; RNA-Seq n= 4, for both groups). Pro-inflammatory cytokine IL-6 is highly expressed at this stage (i), IL-6R, a substrate of Adam17 was also detected with RNA-Seq data. All bar graphs represent Mean ± SEM, \*\**p* < 0.01, \*\*\**p* < 0.001, \*\*\*\**p*< 0.0001. (B) The cytokines *Tnfα* and *Tgfβ1* are highly expressed at this early stage of development, these genes will ultimately influence cell differentiation, inflammation, apoptosis and fibrosis in the adult heart.

## Discussion

To summarise our findings, the inflammatory response, apoptosis and secretion of pro-inflammatory cytokines we identified may play an important role after birth in our extreme mouse model of AC. Macrophage recruitment and activation follow, accompanied by the activation of the fibrotic pathways to facilitate tissue remodelling (Figure 8). Transcriptome data reveal key genes that may drive the inflammatory process and profibrotic cascade which lead to severe cardiomyopathy. The upregulated genes of interest are highlighted at the postnatal (Two weeks) and adult stages (Ten weeks) of the disease. These multiple events are likely to mould the disease phenotype caused by lack of desmoglein 2, a major cadherin and desmosomal protein resulting in disrupted intercalated disc formation, cardiac dysfunction and death.

**Figure 8:**
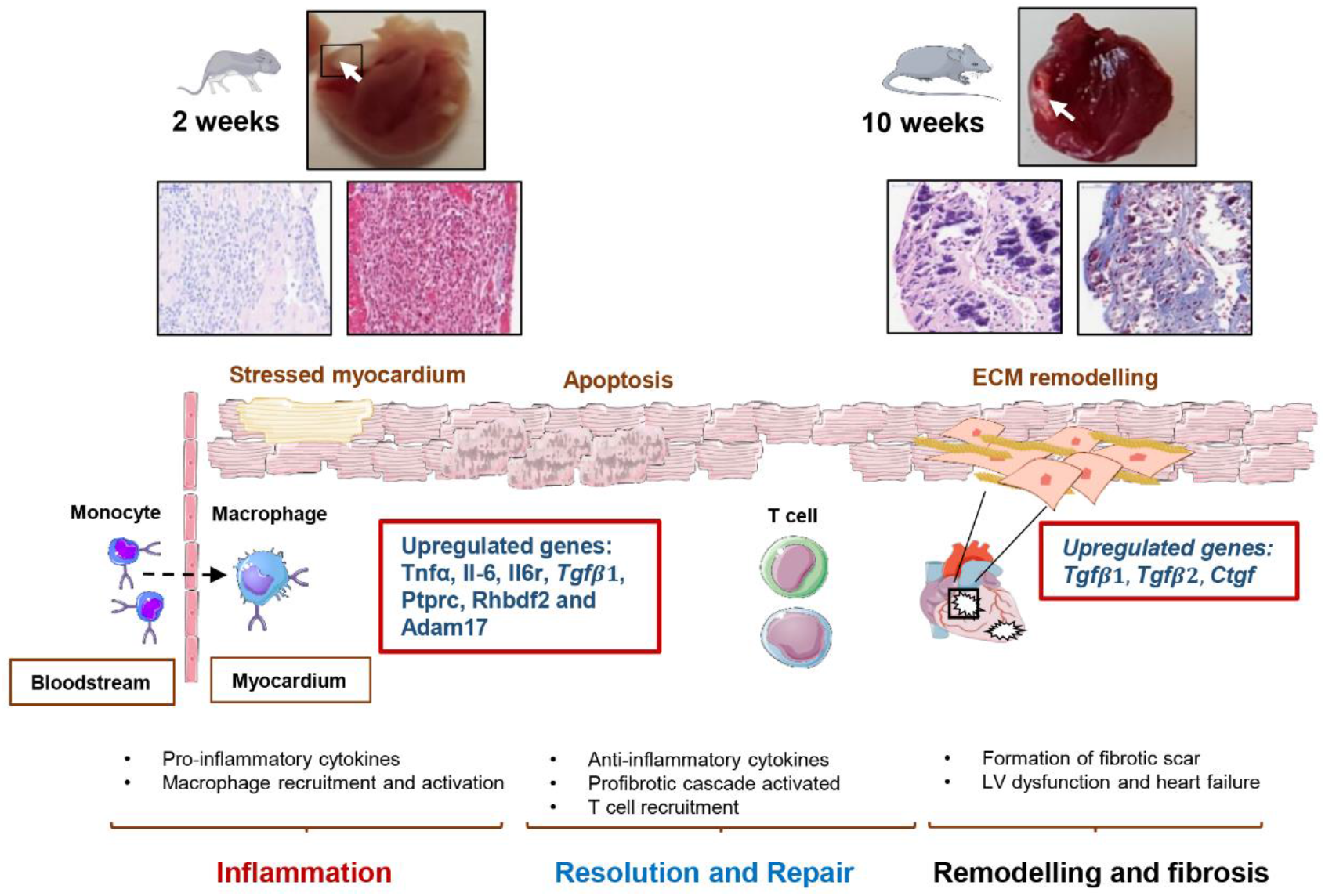
Summary of the proposed key events in cardiac specific murine model of AC. The loss of desmoglein 2 can trigger cell death and initiate the immune response. Postnatal *Dsg2*^−/−^ hearts (Two weeks) suggest inflammation occurs excessively in response to desmosome dysfunction. Our findings reveal key genes are upregulated during the early innate immune response. Pro-fibrotic genes are also expressed at this postnatal period resulting in the pathological condition observed in *Dsg2*^−/−^ adult mice. (This figure was created with templates from Servier Medical Art, licensed under a Creative Common Attribution 3.0 Generic License. http://smart.servier.com).

At two weeks of age, *Dsg2*^−/−^ hearts exhibit myocyte apoptosis, activation of specific inflammatory, stress and profibrotic pathways. In contrast at ten weeks, the Dsg2^−/−^ mouse heart showed severe myocardial dysfunction with extensive fibrosis but a much less prominent immune phenotype. At present, there is little evidence for specific triggers of disease progression for AC and how various mutations affecting the cardiac desmosome can cause such adverse effects. However, it is widely accepted to be worsened by stressors such as exercise and the clinical course can be punctuated by acute arrhythmic episodes (*1*). The potential role of inflammation is only just beginning to be appreciated and may be seen as a potential disease driver rather than a bystander activated as an epiphenomenon (*2, 32*).

The motivation for our study was to examine early inflammation markers in the developing murine heart due to loss of *Dsg2* and provide new insight or markers which may be relevant in the clinical setting. Due to the complexity of the disease, AC diagnosis is often challenging because patients are often asymptomatic until they have suffered a cardiac episode; usually as adults. The disease has been described to display signs of a distinct inflammatory infiltrate in post mortem biopsies from patients with AC (*13*). Recently, six AC paediatric patients were genetically screened and identified with desmosomal mutations (*14*). Cardiac magnetic resonance imaging confirmed myocardial inflammation, these inflammatory events correlated to severe biventricular morphology and reduced cardiac function (*14, 33*). There are phenotypic similarities in these young patients with lesions present in our two week *Dsg2*^−/−^ mouse hearts. The viral trigger was not responsible for myocarditis-like episodes but half of the cases were exercised induced. These myocarditis like episodes or ‘hot phases’ are often associated with a rise in troponin levels and cardiac inflammation.

We have identified some key events which may play a role in the developmental stages of murine AC following birth. In response to cardiac stress and damage, specific macrophage populations remove the dying cardiomyocytes and repair the existing myocardium with fibrosis (*34*). The early immune response at two weeks in *Dsg2*^−/−^ mice was characterised by substantial macrophage infiltration including increased expression of genes associated with pro-inflammatory M1 macrophages (*26, 35*). M1 activated macrophages secrete pro-inflammatory cytokines such as IL-1, IL-6 and TNFα and these and related components in the pathways were increased in expression in our data. In addition to this, stressed cardiomyocytes can release TNFα and TGFβ1 to trigger the inflammatory process and activate the immune cell population to clear apoptotic cells (*36*). Genes linked to M2 macrophages and fibrosis were also increased in expression including TGFβ1. Recent studies have shown that TGFβ1 and CTGF may upregulate IL-11 promoting cardiovascular fibrosis (*30*). It is known that the heart contains a resident population of macrophages and it is thought, that as in other tissues, they have a house-keeping function detecting potential myocardial injury and clearing cellular waste (*37*). In addition, they may also have novel roles such as such as influencing excitability in the atrioventricular node (*38*). The modest electrophysiological phenotype observed in this model is also indicative that arrhythmia may predominantly arise from fibrosis and scar formation rather than pronounced changes in conduction as seen with desmoplakin mutations (*39*). The best characterised injury model is that of myocardial infarction and there is a carefully sequenced process of acute innate inflammation driven largely by macrophages, which ultimately results in repair and scar formation (*34*) (*40, 41*). Whilst there are some similarities, there are also differences. In our case, the two processes of inflammation and repair seem to occur simultaneously but largely terminate at ten weeks. It is unclear if this is due to extensive myocardial damage or whether desmosomal disruption early in life as the heart matures postnatally is the critical driver (*42*).

The inflammatory process that peaks at two weeks is clearly a unique event and is also accompanied by the upregulation of the iRhom2/Adam17 stress pathway and its downstream targets such as Il-6. Our observation that the inflammation is most active early on suggests that intervention with immune supressing drugs would be best given as soon as possible before extensive fibrosis sets in. IL-6 is implicated in chronic inflammation and cancer, and this pathway is one potential therapeutic target for intervention (*43*). There has also been recent interest in IL-11 as a potential target for fibrotic disease (*30*). Glycogen synthase kinase-3 beta (GSK-b), a key player in the pathogenesis of AC can also regulate the inflammatory response (*24*). The potential importance of inflammation in AC has also been indicated by the success of small molecule inhibition of nuclear factor kappa B signalling (*32*) Secondly, any clinical immune signature may only be detected prior to overt disease onset. Conventional treatments such as the use of beta-blockers or implantable cardioverter-defibrillator may benefit from the development of novel diagnostic aids. The prospect of additional studies in murine models at the early stages of cardiac development with known inhibitors could be considered in the future to improve or prevent inflammation and or fibrosis.

## Experimental Procedures

### Generation of genetically modified mice

Animals were cared for according to the Animals (Scientific Procedures) Act 1986. Mice were maintained in an animal core facility under UK Home Office guidelines relating to animal welfare. Mice were kept in individually ventilated, pathogen-free, temperature controlled cages (21-23°C) with 12 hour day/night light cycles and free access to standard rodent chow and water. The *Dsg2* tm1c allele was generated at MRC Harwell (*Oxfordshire, UK*). Our study mice were generated by crossing mice expressing the cardiac αMHC-Cre promoter (Alpha-myosin heavy chain promoter; *Myh6*) with *Dsg2* floxed mice where exon 4-5 of the desmoglein2 gene is flanked by *loxP* sites. This mating resulted in Cre-mediated excision of desmogl—ein 2 in cardiac myocytes, resulting in a frameshift mutation with premature truncation and the product is likely degraded by nonsense-mediated mRNA decay (*24*). Breeding pairs were set up to generate wild type *Dsg2*^+/+^ (Cre^−^, *Dsg2*^flx/flx^) and knockout *Dsg2*^−/−^ mice (Cre^+^, *Dsg2*^flx/flx^). Mice from both sexes were studied at two, six and ten weeks of age to characterise the histology, inflammatory and stress pathways involved and electrophysiological properties of the failing hearts. Ear samples were taken at postnatal day 13 to identify the genotype of the two week cohorts and pups were sacrificed the next day for FACs analysis, RNA sequencing or quantitative RT-PCR analysis.

### In vivo assessment of cardiac function

*In-vivo* cardiac assessment was performed with VisualSonics (*VisualSonics Inc., Toronto, Canada*) Vevo3100 high-resolution ultrasound scanner with a 30-MHz Frequency transducer. Mice were anaesthetised with 1.5 % (v/v) isoflurane whilst body temperature was maintained at 37 °C with a temperature probe. B and M mode parasternal long-axis and short-axis views were acquired to evaluate cardiovascular structures and cardiac function. Doppler scans were also taken to determine blood flow velocity and direction, as evidenced by colour differential. ECGs were also obtained with the built-in ECG electrode contact pads. Measurements were taken at two time points to assess cardiac function at six weeks (in juvenile mice) and later at ten weeks (adult mice) when the disease in the *Dsg2*^−/−^ mouse should manifest (n=8, all groups). ECG measurements were processed in LabChart (*ADI Instruments, UK*) and data was analysed in Prism 8.0. The Vevo LAB 3.1.1 analysis software (*VisualSonics Inc., Toronto, Canada*) was used for quantifying and characterising our mouse model of AC. The most commonly used variables to evaluate systolic function are fractional shortening (FS) and ejection fraction (EF) measured from changes in chamber dimensions during systole (S) and diastole (D). Measurements and calculations were obtained from short axis M mode with the Left ventricle (LV) trace method; results were saved and processed in Excel before statistical analysis in Prism 8.0.

### Electrophysiological Mapping

We used a flexible multielectrode array (*FlexMEA; Multichannel systems, Germany*) to study the electrophysiological properties of the intact heart as we have previously described (*22, 23*). The data collected was used to calculate conduction velocity (CV), mean increase in delay (MID) and effective refractory period (ERP) from our study cohort (n=8, both groups). Hearts were perfused in the Langendorff mode with normal oxygenated Krebs solution (Ca^2+^ 1.4mM) at 16.5 ml/min. A unipolar silver chloride stimulation electrode and flexible 32-pole multi-electrode array (MEA) (*FlexMEA, Multielectrode System, Germanys*) was placed on the ventricular epicardium and an S_1_S_2_ decremental protocol was performed to measure the ventricular effective refractory period (VERP). A biphasic pulse of amplitude 2 volts and duration 0.5 ms was used for stimulation, with S_1_S_2_ intervals reduced from 150 ms by decrements of 5 ms to 100 ms followed by decrements of 2 ms until tissue refractoriness was reached. Arrhythmogenicity was further tested for by applying stimulating trains of 100 beats at coupling intervals progressively reduced from 100 ms. Ventricular tachycardia was defined as a ventricular arrhythmia lasting more than 2 seconds.

All analysis of murine electrophysiology was performed using custom software running in Matlab, v2014b (*The Mathworks Inc., MA, USA*). The time point of local activation was taken at the steepest negative gradient of the unipolar electrogram. Conduction velocities were determined using a gradient method, with conduction velocity defined as the inverse of the gradient in activation times across the array. Electrodes with significant noise were excluded, and all electrograms and time points were checked manually. MID, a well-validated measure of inducibility of conduction delay, was calculated by determining the area under the conduction-delay curve. The mean timing of the activation time of all recording electrodes was used for each measurement of conduction delay, and the MID defined as the unit increase in conduction delay per unit reduction in S1S2 coupling interval (ms/ms). Stimulation protocols were performed in normal Krebs Henseleit solution (*Sigma, UK*).

### Tissue collection and morphometric analysis

Following the *ex-vivo* whole heart experiments, hearts were rinsed immediately in PBS before they were placed into a tissue holder and cut longitudinally using both right and left atrial appendages as a guideline. Excess moisture from both halves were blotted off before the weight was of the individual hearts were recorded. One half was fixed in 10% formalin for 24 hours and paraffin-embedded for histology and immunostaining. The other half was cut and approximately 30 mg of the left ventricle was stored in RNAlater (*Sigma, UK*). Samples were obtained from two week and ten week old mice for qPCR analysis (n= 5, both groups). Whole hearts were collected from a separate cohort of two week old mice for FACs analysis (n= 7, both groups).

### Murine histology and immunohistochemistry

Mouse hearts collected from two and ten week old animals were rinsed thoroughly in PBS to remove excess blood and fixed in 10 % formalin for at least 24 hours. After the fixation process, they were washed twice in PBS and stored in 70 % ethanol before paraffin embedding. Paraffin-embedded myocardium were cut at 5 μm thick sections and mounted on clear, plus microscope slides. For histological analysis, sections were stained for haematoxylin and eosin with automated Leica autostainer XL system (*Leica Biosystems, UK*) and Trichrome stain kit (*ab150686, Abcam, UK*) to detect cardiac fibrosis. (according to manufacturer’s instructions).

### DAB Staining

To identify immune cell populations from two week *Dsg2*^−/−^ hearts, we employed the chromogenic tissue staining method using the DAB detection system. The automated Ventana Classic and XT system (*Roche Diagnostics, UK*) was used to process our paraffin embedded heart samples. The antibodies used were: rat monoclonal CD45 (*ab25386, Abcam, UK*) and rat monoclonal F4/80 *(MCA497GA Clone CI: A3-1 Serotec, UK*). The OmniMap anti rabbit HRP kit (*760-4311, Roche Diagnostics, UK*) was used to detect the antibodies of interest. The samples were counterstained with haematoxylin; slides were sealed with mountant and coverslip and allowed to dry. Sections from *Dsg2*^+/+^ and *Dsg2*^−/−^ hearts were also stained with *Dsg2* antibody (*Ab150372, Santa Cruz UK)* to confirm *Dsg2* was deleted in the heart. Slides from the various histological stains were scanned with the Pannoramic 250 High Throughput Scanner (*HistoTech, Budapest, Hungary*) and representative images from two week and ten week samples with scale bars were processed with Panoramic Viewer software (*HistoTech, Budapest, Hungary*).

### TUNEL Staining

Paraffin samples from two week old mice were stained for TUNEL assay to confirm apoptosis caused by DNA Fragmentation. The ApopTag^®^ Plus In Situ Apoptosis Fluorescein Detection Kit (*S7111 Sigma-Aldrich, UK*) was used following manufacturer’s instructions. The slides were kept in the dark and imaged with confocal microscopy (*Zeiss LSM 880 with Airyscan Fast, Carl Zeiss, UK*). The number of TUNEL positive cells and total number of cell nuclei counterstained with DAPI were calculated in Image J with ITCN plugin (Nuclei counter) to generate the % of TUNEL positive cells (n= 3 both groups).

### RNA sequencing analysis (RNA-Seq)

RNA samples from two and ten week hearts (n= 4, both groups) were assessed for quantity and integrity using the NanoDrop 8000 spectrophotometer V2.0 (*ThermoScientific, USA*) and Agilent 2100 Bioanalyser (*Agilent Technologies, Waldbronn, Germany*). All samples displayed low levels of degradation with RNA integrity numbers (RIN) between 7.6 and 9.1. One hundred ng of total RNA from each sample was used to prepare total RNA libraries using the KAPA Stranded RNA-Seq Kit with RiboErase (*KAPA Biosystems, Massachusetts, USA*). Fragmentation prior to first strand cDNA synthesis was carried out using incubation conditions recommended by the manufacturer for intact RNA samples (94 °C for 6 minutes (min)), and 14 cycles of PCR were performed for final library amplification. Resulting libraries were quantified using the Qubit 2.0 spectrophotometer *(Life Technologies, California, USA*) and average fragment size assessed using the Agilent 2200 Tapestation (*Agilent Technologies, Waldbronn, Germany*). Equimolar amounts of each sample library were pooled together, and 75 bp paired-end reads were generated for each library using the Illumina NextSeq®500 High-output sequencer (*Illumina Inc., Cambridge, UK*).

In Galaxy v19.05, FASTq files were mapped to GRCm38.97 mouse ensemble genome using RNAstar v2.6, gene and transcript counts made with StringTie v1.3.4, GTF merged tables of all samples were made by StringTie merge v1.3.4, recounted using StringTie and differential expression tested by edgeR v3.24.1 / Deseq2 v1.18.1. KEGG Pathway analyses were generated using DAVID Bioinformatics Resources, version 6.8 (https://david.ncifcrf.gov/home.jsp) (*44, 45*). Gene lists for analysis were identified using a cut off of adjusted *p*-value < 0.05 and log2 fold-change ≥ 1 and ≤ −1. KEGG outputs were filtered using false discovery rate (FDR) *p*-value < 0.05, and data was log10 transformed for visualisation. Immune signatures were selected for the two week samples. The heatmap was generated from 41 differentially expressed immune-related genes. An adapted version (including several macrophage-related genes) of the ‘Mouse nCounter^®^ Immunology Panel’ (nanoString) was used to filter genes, and differentially expressed genes were defined as adjusted *p*-value < 0.05, log2 fold-change ≥ 1.5 and ≤ −1.5, base mean > 500. Heatmap was plotted using the pheatmap package (v1.0.12) in R (v3.6.2). RNA-Seq data are presented with FPKM (Fragments Per Kilobase of transcript per Million mapped reads) values.

The data discussed in this publication have been deposited in NCBI's Gene Expression Omnibus (*46*) and are accessible through GEO Series accession number GSE153124 (https://www.ncbi.nlm.nih.gov/geo/query/acc.cgi?acc=GSE153124).

### Quantitative real-time PCR (qRT-PCR)

Total RNA (from ~30 mg of tissue) was extracted from two and ten week mouse heart samples stored in RNA Later with the RNeasy fibrous tissue kit (*74704, Qiagen, UK*). RNA was quantified using a NanoDrop spectrophotometer (*Thermo Fisher Scientific, UK*) and total RNA was DNase I treated and reverse-transcribed using the High-capacity cDNA reverse transcription kit (*4368813, Applied Biosystems, Life Technologies, UK*). Fifty ng of cDNA was used for qRT-PCR, which was performed using customised Taqman gene expression assays (*Applied Biosystems, UK*). Commercially available probes for genes of interest were used (mouse gene names in italics): Desmoglein 2 *Dsg2*, CD45 *Ptprc,* interleukin-6 *Il-6,* inactive rhomboid protein 2 *Rhbdf2,* disintegrin and metalloproteinase domain-containing protein 17 *Adam17,* collagen type I alpha 1 chain *Col1A1*, collagen type 3 alpha 1 chain *Col3A1*, connective tissue growth factor *Ctgf*, transforming growth factor β1 *Tgfβ1* and transforming growth factor β2 *Tgfβ2* (Table 2). Each gene was assayed in triplicate and relative expression was calculated by using the comparative CT method normalised to GAPDH (n= 5 for both groups and timepoints). The data are presented as relative changes when compared the control group (*Dsg2*^+/+^) at two weeks (set at one arbitrary units).

**Table 2.**
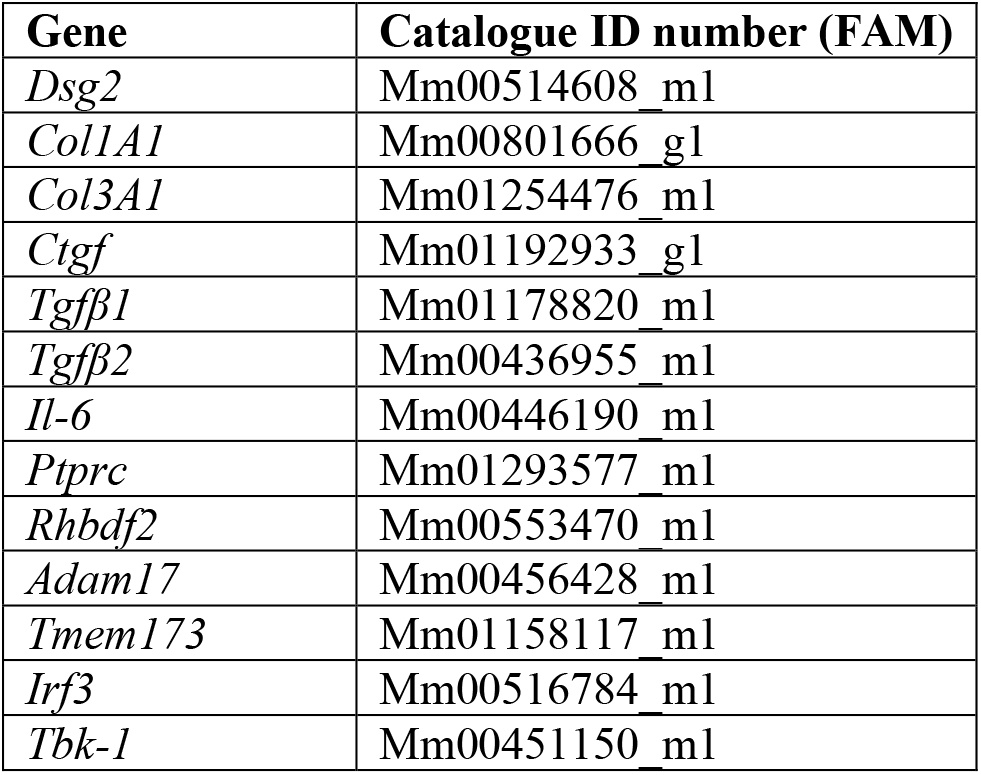
List of mouse probes used in this study. All probes were obtained from Thermofisher Scientific UK.

### Fluorescence-activated cell sorting (FACS)

Mice were sacrificed by neck dislocation, hearts were harvested and rinsed in PBS. The tissue was cut into small pieces and then processed in digestion buffer containing HANKS Medium (10-527F, Lonza, UK), 608 U/ml of Collagenase I (C9891m Sigma, UK), 187.5 U/ml Collagenase XI (C7657, Sigma, UK), 90 U/ml Hyaluronidase (H3884, Sigma, UK), 90 U/ml DNAse (D4263, Sigma, UK). Samples were incubated at 37 °C in a shaking incubator at 100 rpm for 1.5 hours. The contents were then passed through 70 μm cell strainers to remove debris and the filtrate diluted with PBS. These were centrifuged to create a pellet and the cells resuspended at a concentration of 2×106 cells/50 μl.

Fc block was performed using TruStain FcX™ CD16/32 antibody (anti-mouse) to avoid non-specific binding (101320 Biolegend, UK) according to the supplier protocols The fluorochrome-conjugated antibodies used in the panel were obtained from Biolegend, UK: CD11c-BV605 (117333), CD8-AF488 (100723), Ly6G-PE (127607), F4/80-PE/Cy5 (123111), CD11b-PE/Cy7 (101215), CD4-APC (100412), CD45.2-AF700 (109822). The antibodies Ly6C -eF450 (48-5932-80) and CD3-PerCP/eF710 (46-0032-80) were obtained from eBioscience, UK. Fluorescence minus one (FMO) controls were included for each marker. The samples were analysed using a LSR Fortessa system (BD bioscience, UK) equipped with BD FACS DivaTM software (BD Biosciences, UK), followed by analysis using FlowJo (Version 10.2,Treestar, Ashland, Oregon). The CD45+ counts were normalised to the total number of events run, the final data show the percentage of the different cell types that were CD45+.

### Statistical analysis

All results are presented as mean + SEM where n is the number of mice used. Statistical analyses were conducted using GraphPad Prism (*version 8.0; GraphPad software, California, USA*). For comparison of two sets of data, a two-tailed, unpaired Student’s t-test was used. For the analysis of three or more groups, one-way ANOVA followed by Dunnett’s or Tukey multiple comparisons tests were used where *p*<0.05 was statistically significant.

## Acknowledgments

We would like to thank the Queen Mary University of London (QMUL) core services, in particular Barts and the London Genome Centre and Bart Cancer Institute (BCI) Pathology services for their assistance in processing our samples. We are grateful to Professor Mary Sheppard at St George’s Hospital, London for discussions. We acknowledge Reya Srivastava for her assistance genotyping our study mice. Our work was supported by the British Heart Foundation grant RG/13/19/30568 ‘’Unravelling the molecular and mechanistic complexity of ARVC via the skin”.

## Conflicts of interest

The authors declare that they have no conflicts of interest with the contents of this article.

## Author contributions

K.E. Ng, D.P. Kelsell and A. Tinker designed the research conceptual framework of the study. K.E. Ng, P.J. Delaney and D.J. Thenet generated the transgenic mice, performed experiments and analysed data. E. Tsisanova completed TUNEL experiments. S.M Walker assisted with FACs analysis. K.E. Ng, S. Murtough, C.M. Webb and D.J Pennington analysed the RNA-Seq data. R. Pink provided the training and expertise for RNA-Seq analysis. J.D. Westaby provided expertise and aided with histological analysis. K.E. Ng, D.P. Kelsell and A. Tinker wrote the manuscript.

---

**Figure S1:**
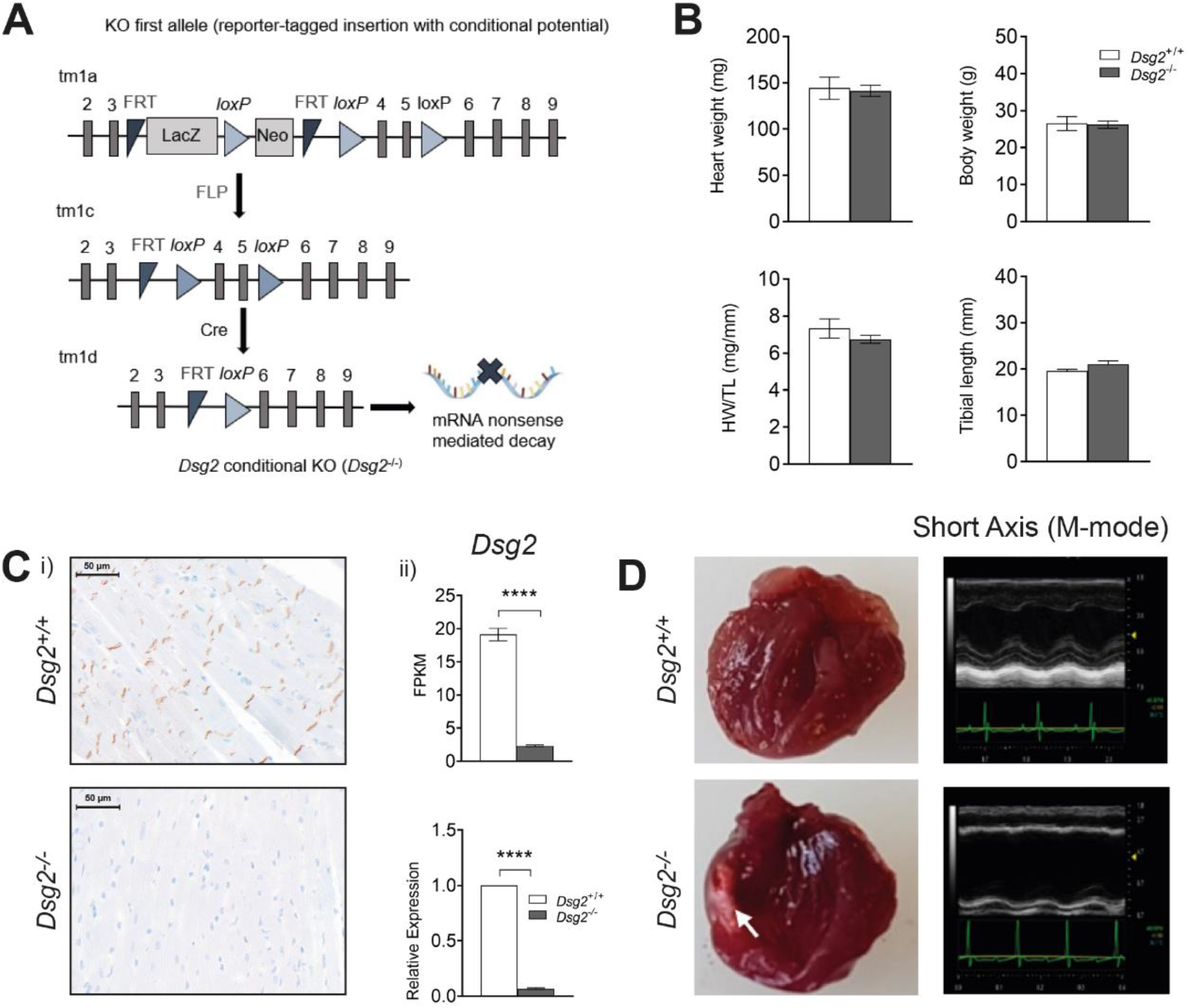
(A) Cre/LoxP targeting strategy for the deletion of exon 4-5 of *Dsg2*. Schematic depicting the targeting vector construct with recombined *Dsg2* locus (tm1a). The *Dsg2* (^+/flx^) allele after Flp-mediated excision (tm1c). Tissue-specific deletion of *dsg2* with Alpha MHC Cre recombinase. (B) Morphometric analysis of *Dsg2*^+/+^ and *Dsg2*^−/−^ at ten weeks of age, we measured body weights (g), tibial length (mm), whole heart weights (mg). Heart weights were normalised to tibial length (mg/mm) (n= 8, for both groups). There were no significant differences between *Dsg2*^+/+^ and *Dsg2*^−/−^ adult samples.(C) i) DAB immunohistochemistry confirms the absence of *Dsg2* in the *Dsg2*^−/−^ heart, note in the *Dsg2*^+/+^ heart, *Dsg2* resides at the intercalated disc. Separate cohorts of mice were used for ii) RNA-Seq (n= 4, for both groups) and qRT-PCR (n= 5, both groups) to validate deletion of *Dsg2*. Data represent Mean ± SEM, \*\*\*\**p*< 0.0001. (D) Cardiac assessments for adult mice (n= 8, for both groups). In all our subjects, the left ventricle in the *Dsg2*^−/−^ heart was dilated with little contractility when compared to *Dsg2*^+/+^ heart. The gross morphology of the *Dsg2*^−/−^ heart confirmed the walls of the ventricles were considerably thinner and white fibrotic plaques were present.

**Figure S2:**
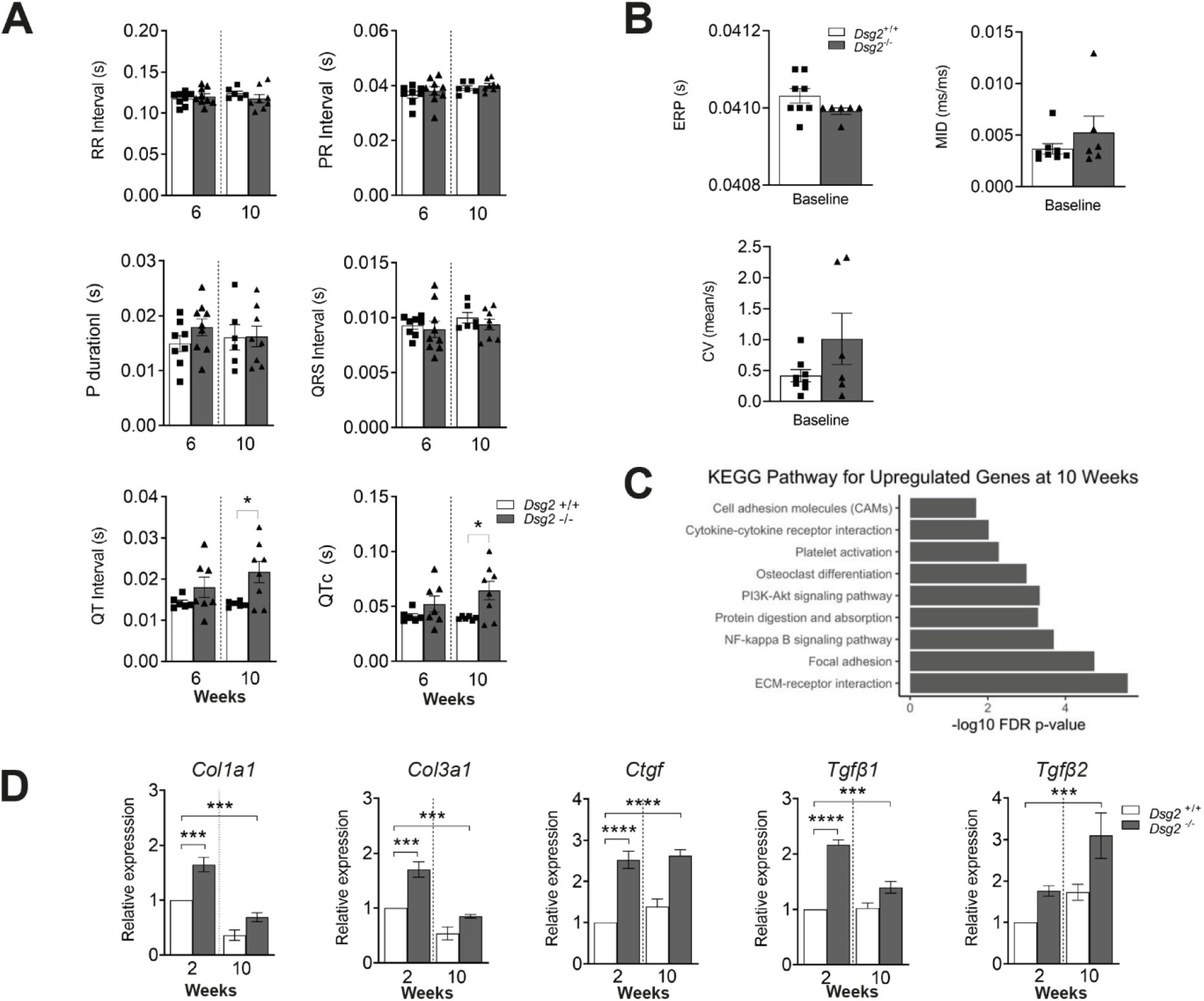
(A) Bar graph with scatterplots to show data for ECG parameters such as RR (s) and QRS interval (s) obtained from ECG recordings from mice undergoing cardiac assessment (echocardiography) at six weeks and ten weeks of age. QT (s) data is significant at the ten weeks between the *Dsg2*^+/+^ and *Dsg2*^−/−^ (n= 8, for all groups). Data represent Mean ± SEM. \**p* < 0.05. (B) Baseline electrophysiological measurements (ERP, CV and MID) acquired from Flex MEA array (*Dsg2*^+/+^ n= 8; *Dsg2*^−/−^ n= 6). (C) KEGG full pathway output for upregulated genes at ten weeks. (C) KEGG full pathway output for upregulated genes at ten weeks. (D) qRT- PCR results for fibrosis markers *Col1a1*, *Col3a1*, *Ctgf*, *Tgfβ1* and *Tgfβ2* at two and ten weeks (n= 5, all groups). Fibrosis is also an early event and occurs when the immune response is triggered at two weeks in *Dsg2*^−/−^ hearts. Bar graphs show Mean ± SEM \*\*\**p* < 0.001, \*\*\*\**p* < 0.0001.

